# Learning the statistics of pain: computational and neural mechanisms

**DOI:** 10.1101/2021.10.21.465270

**Authors:** Flavia Mancini, Suyi Zhang, Ben Seymour

## Abstract

Pain invariably changes over time, and these temporal fluctuations are riddled with uncertainty about body safety. In theory, statistical regularities of pain through time contain useful information that can be learned, allowing the brain to generate expectations and inform behaviour. To investigate this, we exposed healthy participants to probabilistic sequences of low and high-intensity electrical stimuli to the left hand, containing sudden changes in stimulus frequencies. We demonstrate that humans can learn to extract these regularities, and explicitly predict the likelihood of forthcoming pain intensities in a manner consistent with optimal Bayesian models with dynamic update of beliefs. We studied brain activity using functional MRI whilst subjects performed the task, which allowed us to dissect the underlying neural correlates of these statistical inferences from their uncertainty and update. We found that the inferred frequency (posterior probability) of high intensity pain correlated with activity in bilateral sensorimotor cortex, secondary somatosensory cortex and right caudate. The uncertainty of statistical inferences of pain was encoded in the right superior parietal cortex. An intrinsic part of this hierarchical Bayesian model is the way that unexpected changes in frequency lead to shift beliefs and update the internal model. This is reflected by the KL divergence between consecutive posterior distributions and associated with brain responses in the premotor cortex, dorsolateral prefrontal cortex, and posterior parietal cortex. In conclusion, this study extends what is conventionally considered a sensory pain pathway dedicated to process pain intensity, to include the generation of Bayesian internal models of temporal statistics of pain intensity levels in sensorimotor regions, which are updated dynamically through the engagement of premotor, prefrontal and parietal regions.

## INTRODUCTION

In recent years, our understanding of pain has shifted from viewing it as a simple responsive system to a complex predictive system, that interprets incoming inputs based on past experience and future goals (Fields, 2018). Indeed, all types of pain response, including perception, judgement and decision-making, are invariably and often strongly shaped by what pain is being predicted, and the nature of this influence gives clues regarding the fundamental architecture of the pain system in the brain (Büchel et al., 2014; Seymour and Mancini, 2020; Roy et al., 2014; Wiech, 2016). To date, most experimental strategies to study prediction have come from explicit cue-based paradigms, in which a learned or given cue, such as visual image, contains the relevant information about an upcoming pain stimulus. (Atlas et al., 2010; Büchel et al., 2014; Fazeli and Büchel, 2018; Geuter et al., 2017; Zhang et al., 2016). However, a much more general route to generate predictions relates to the background statistics of pain over time - the underlying base-rate of getting pain, and of different pain intensities, at any one moment. In principle, the pain system should be able to generate predictions based on how pain changes over time, in absence of external cues. This possibility is suggested by research in other sensory domains, showing that the temporal statistics of sequences of inputs are learned and inferred through experience - a process termed temporal statistical learning (Dehaene et al., 2015; Frost et al., 2015; Fiser and Aslin, 2002; Kourtzi and Welchman, 2019; Turk-Browne et al., 2005; Wang et al., 2017). We hypothesise that temporal statistical learning also occurs in the pain system, allowing the brain to infer the prospective likelihood of pain by keeping track of ongoing temporal statistics and patterns. In this way, pain should effectively act as the cue for itself, instead of utilising a cue from a different sensory modality. This may be especially important in clinical contexts, in which pain typically comes in streams of inputs changing over time (Kajander and Bennett, 1992).

Here, we tested this hypothesis by designing a frequency learning paradigm involving long, probabilistic sequences of noxious stimuli of two intensities (low and high) that could suddenly change. We tested people’s ability to generate explicit predictions about the probability of forthcoming pain, and probed the underlying neural mechanisms. In particular, following evidence in other sensory domains (Meyniel et al., 2016), we proposed that the brain uses a optimal Bayesian strategy to infer the background temporal statistics of pain. Importantly, this approach may allow us to map core regions of the pain system to specific functional information processing operations: the temporal prediction of pain, its uncertainty and update. Our hypothesis predicts that the predictive inference of pain stimuli should be encoded largely within pain processing brain regions (Conway and Christiansen, 2005). The uncertainty of the prediction is expected to implicate multisensory, intraparietal regions, as shown previously using visual and auditory stimuli (Meyniel and Dehaene, 2017).

## RESULTS

Thirty-five participants (17 females; mean age 27.4 years old; age range 18-45 years) completed an experiment with concurrent brain fMRI scanning. They received continuous sequences of low and high intensity painful electrical stimuli, wherein they were required to intermittently judge the likelihood that the next stimulus was of high versus low intensity (figure 2 a). We designed the task such that the statistics of the sequence could occasionally and suddenly change, which meant that the the sequences effectively incorporated sub-sequences of stimuli. The statistics themselves incorporated two types of information. First, they varied in terms of the relative frequency of high and low intensity stimuli, to test the primary hypothesis that frequency statistics can be learned. Second, sequences also contained an additional aspect of predictability, in which the conditional probability of a stimulus depended on the identity of the previous stimulus (i.e. its transition probability). By having different transition probabilities between high and low stimuli within subsequences, it is possible to make a more accurate prediction of a forthcoming stimulus intensity over-and-above simply learning the general background statistics. For instance, if low pain tends to predict low pain, and high predicts high, then one tends to get ‘clumping’ patterns of pain (runs of high or low stimuli); or conversely if high predicts low and vice versa, one tends to get alternating patterns. Both might have the same overall frequency of high and low pain, but better predictions can be made by learning the *temporal patterns* within. Thus we were able to test the supplementary hypothesis that humans can learn the specific transition probabilities between different intensities, as shown previously with visual stimuli (Meyniel et al., 2016).

**Figure 1.**
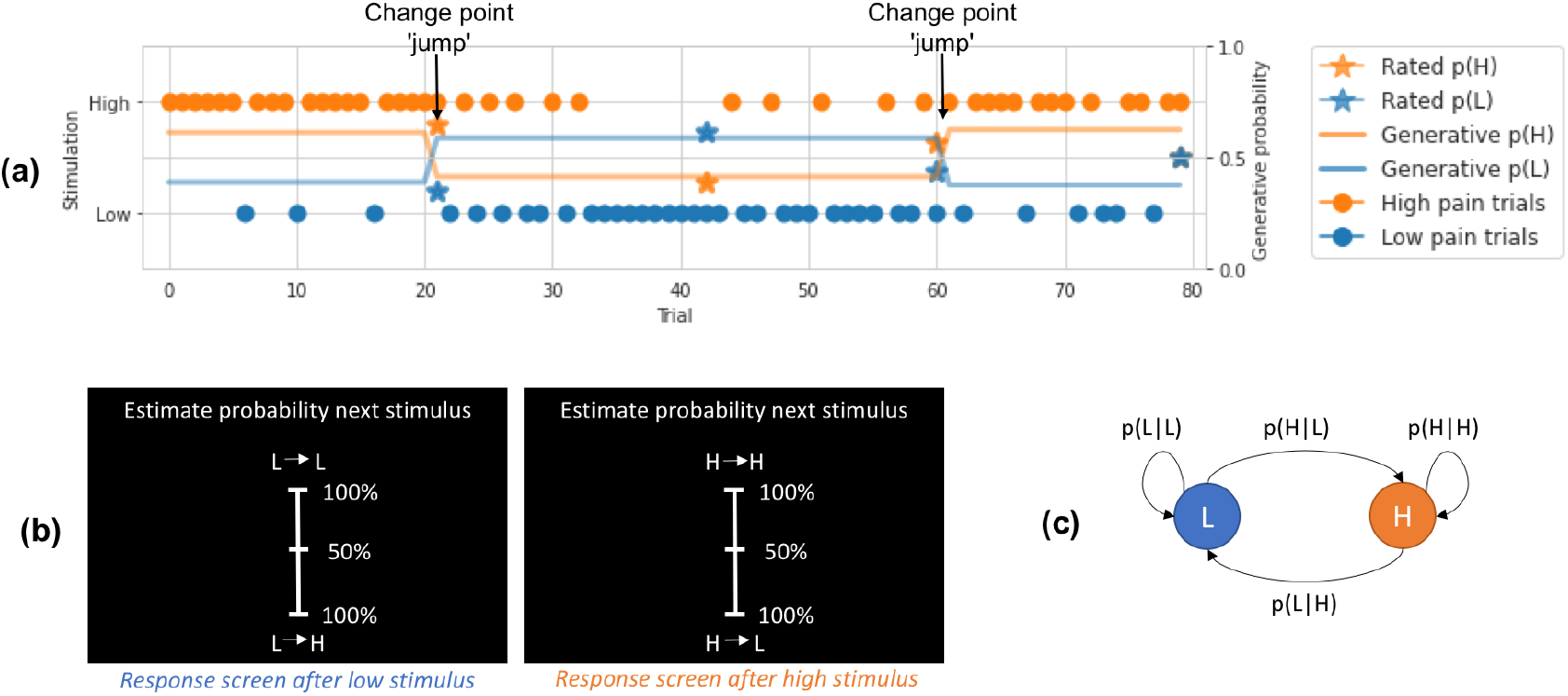
Behavioural task and model explanation. (a) Example trials from a representative participant, showing the true probability of high (H) and low (L) stimuli given current stimuli, trial stimulation given, and participant rated probabilities. Arrows pointing to jump points of true probabilities, where a large change happens. (b) Participant rating screens during the task, where they were asked to estimate the identity of the upcoming stimulus given the current one. For example, after a low stimulus participants would be asked to rate the probability of the upcoming stimulus being low (L -> L) or high (L -> H). (c) Markovian generative process of the sequence of low and high intensity stimuli, depicted in a. The transition probability matrix was resampled at change points, determined by a fixed probability of a jump.

**Figure 2.**
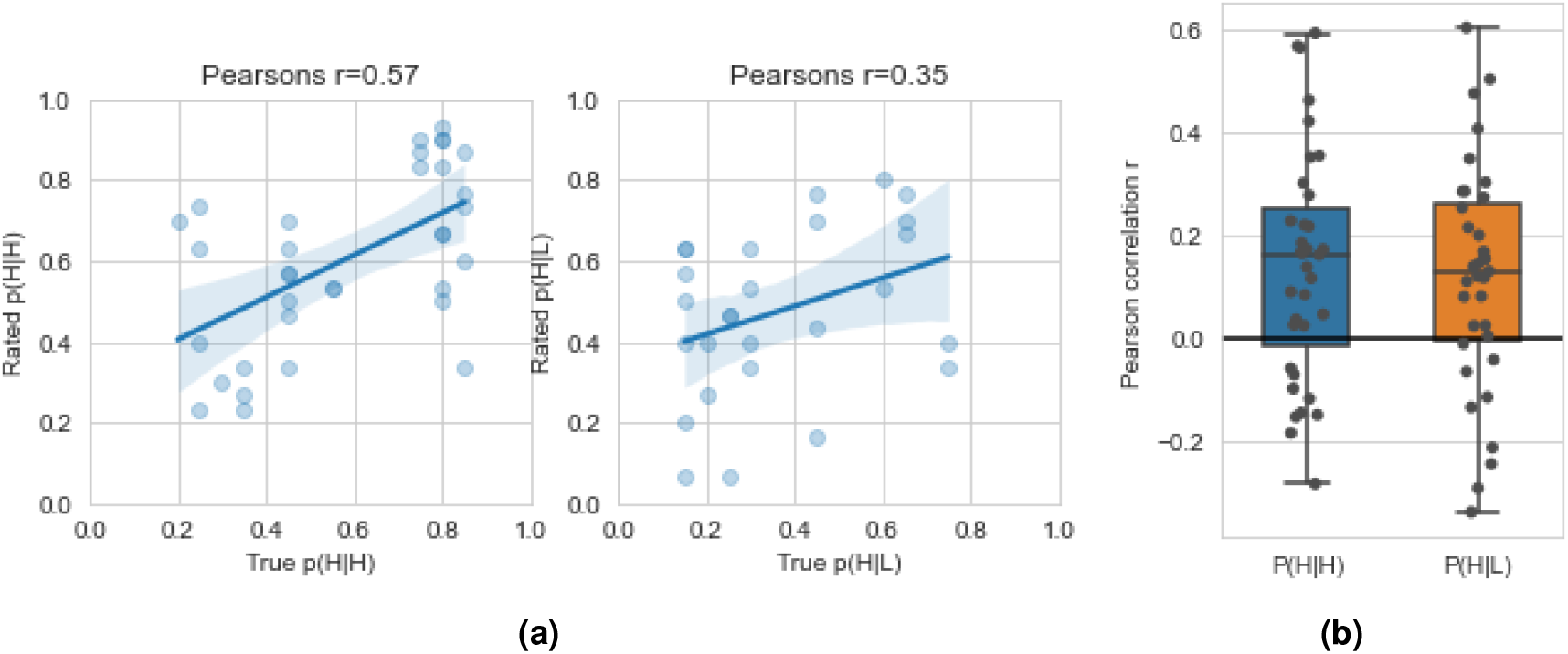
Behavioural results. (a) True vs rated probabilities for p(H|H) and p(H|L) from an example participant, a positive correlation suggests the participant correctly learned the stimuli probability, (b) Pearson’s r for true vs rated probabilities for p(H|H) and p(H|L) within individual participants.

At the beginning of the experiment, participants were informed that the sequence was set by the computer and could occasionally change at any point in time. This design mirrored a well-studied task used to probe statistical learning with visual stimuli (Meyniel and Dehaene, 2017); participants were explicitly and occasionally asked to estimate the probability of forthcoming stimuli (figure 2 b). The sequence was thus defined by a set of transition probabilities: the probability of high or low pain following a high pain stimulus; and the probability of high or low pain following a low pain stimulus (i.e. a Markovian transition matrix; see example in figure 2 c). Occasionally, these probabilities were suddenly resampled, such that in fact the total task length of 1300 stimuli (split into 5 blocks) comprised typically about 50 subsequences (mean 25 ± 4 stimuli per subsequence). Participants were not explicitly informed when these changes happened. Within these subsequences, the frequency of high (versus low) stimuli varied from 15% to 85%, and figure 2 a illustrates an example of a snapshot of a typical sequence, showing a couple of ‘jump’ points where the probabilities change. Figure 2 b shows the rating screen, with ratings being required on 4.8% of stimuli. Before the main experimental scanning session, subjects practiced the task for an average of roughly 1200 trials before the MRI sessions.

### Behavioural results

Participants were able to successfully learn to predict the intensity (high versus low) of the upcoming painful stimulus within the sequence. Fig 2a shows the positive correlation between stimulus rated and true probabilities for low and high pain respectively for an example individual (Pearson correlation for this participant p(H|H) r=0.567, p=4.61e-4, p(H|L) r=0.348, p=0.075; see supplementary figs 1-2 for plots from all subjects). Across subjects, the within-individual Pearson’s r between true and rated probabilities was significantly above zero. (Fig 2b, 26 out of 35 subjects had r>0: p(H|H) r=0.138 ± 0.225, t(34)=3.65, p=0.00088, Cohen’s d=0.871; p(H|L) r=0.117 ± 0.220, t(34)=3.15, p=0.0034, Cohen’s d=0.752; see also supplementary figures 1-2; note that p(H|L) and p(L|L) are reciprocal, as well as p(H|L) and p(L|L)).

### Behavioural data modelling

#### Model choice

Based on previous evidence in other sensory domains, we hypothesised that subjects use an optimal Bayesian strategy to infer the statistics over time (Meyniel, 2020; Meyniel et al., 2016). We fit subjects’ ratings to four variations of a Bayesian model, according to two factors: first, sequence inference through stimulus frequency (by assuming the sequence as generated by a Bernoulli process, where subjects track how often they encountered previous stimuli), versus inference through transition probability (by assuming the sequence follows a Markov transition probability between successive stimuli, where the subject tracks such transition of previous stimuli). This distinguishes between whether participants learn simple statistics (our primary hypothesis), or are able to learn the full transition probabilities (supplementary hypothesis). The second factor was to whether the model incorporates the possibility of sudden changes (jumps) in stimuli probability, as occurs in the task paradigm, or ignores such possbilities (fixed). To compare against alternative models, we also fit a basic reinforcement learning model (Rescorla-Wagner with fixed learning rate, which is an established model of Pavlovian conditioning; (Rescorla et al., 1972)) and a baseline random model that assumes constant probabilities throughout the experiment for high and low pain respectively.

#### Model fitting

The selected models estimate the probability of a pain stimulus’ identity in each trial. The values predicted by the model can be fitted to actual subject predictive ratings gathered during the experiment. A model is considered a good fit to the data if the total difference between the model predicted values and the subjects’ predictions is small. Within each model, free parameters were allowed to differ for individual subjects in order to minimise prediction differences. For Bayesian ‘jump’ models, the free parameter is the prior probability of sequence jump occurrence. For Bayesian fixed models, the free parameters are the window length for stimuli history tracking, and an exponential decay parameter that discounts increasingly distant previous stimuli. The RL model’s free parameter is the initial learning rate, and random model assumes a fixed high pain probability that varies across subjects. The model fitting procedure minimises each subject’s negative log likelihood for each model, based on residuals from a linear model that predicts subject’s ratings using learning model predictors. The smaller the sum residual, the better fit a model’s predictions are to the subject’s ratings.

#### Model comparison

We compared the different models using the likelihood calculated during fitting as model evidence. Fig 3a showed model frequency, model exceedance probability, and protected exceedance probability for each model, fitted for fMRI sessions of the experiment. Both comparisons showed the winning model was the ‘Bayesian jump frequency’ model inferring both the frequency of pain states and their volatility, producing predictions significantly better than alternative models (Bayesian jump frequency model frequency=0.563, exceedance probability=0.923, protected exceedance=0.924). Fig 3b reports the model evidence for each subject; it shows that, although the majority (n=23) of participants were best fit by the model that infers the background frequency, some participants (n=12) were better fit by the more sophisticated model that infers specific transition probabilities.

**Figure 3.**
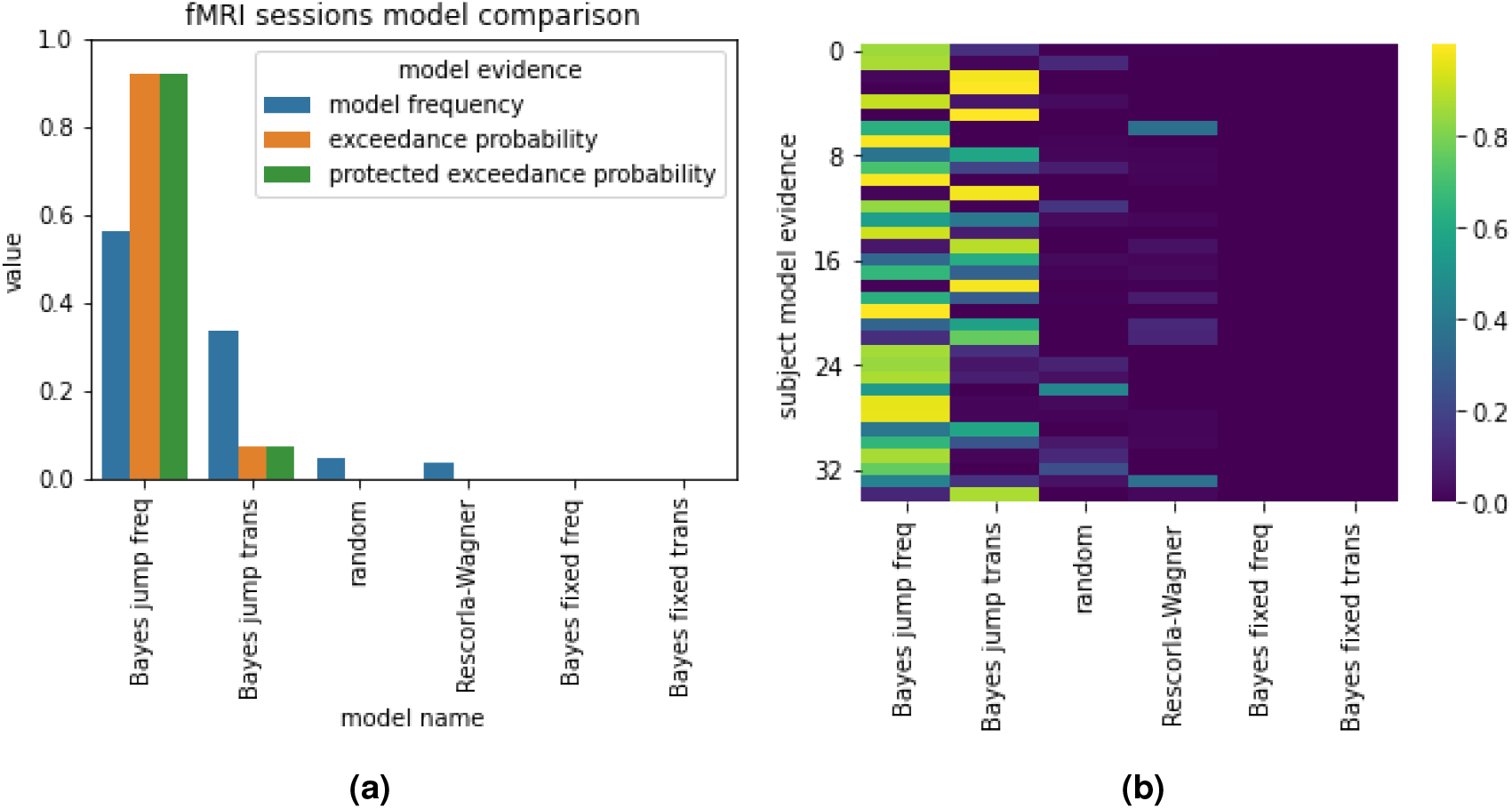
Model comparison results. (a) Bayesian model comparison based on model fitting evidence, in fMRI sessions. Subjects’ predictive ratings of next trial’s pain intensity were fitted with posterior means from Bayesian models, values from Rescorla-Wagner (reinforcement learning) model, and random fixed probabilities. Bayesian jump frequency model (assuming jumps in sequence and inference with stimuli frequency) was the winning model in both cases. (b) Individual subject model evidence.

### Neuroimaging results

We used the winning computational model to generate trial-by-trial regressors for the neuroimaging analyses. The rationale of this approach is that neural correlation of core computational components of a specific model provides evidence that and how the model is implemented in the brain (Cohen et al., 2017). First, a simple high*>*low pain contrast identified BOLD responses in the right thalamus, sensorimotor, premotor and supplementary motor cortex, insula, anterior cingulate cortex and left cerebellum (with peaks in laminae V-VI), consistent with the known neuroanatomy of pain responses (fig 4, table 1). The opposite contrast (low>high pain) is reported in Supplementary Figure 3 and Supplementary Table 1.

**Table 1.** High pain > low pain stimuli activation clusters (FWE p<0.05).

|  | Cluster ID | X | Y | Z | Peak Stat | Cluster Size (mm3) |
| --- | --- | --- | --- | --- | --- | --- |
| 0 | 1 | 13 | -17 | 5 | 6.886 | 2587 |
| 1 | 1a | 13 | -22 | -3 | 4.957 |  |
| 2 | 2 | 33 | -22 | 58 | 6.524 | 5247 |
| 3 | 2a | 30 | -19 | 49 | 5.953 |  |
| 4 | 2b | 16 | -17 | 68 | 5.524 |  |
| 5 | 3 | 37 | -17 | 15 | 5.742 | 1186 |
| 6 | 4 | 6 | -7 | 49 | 5.128 | 3486 |
| 7 | 4a | 0 | -2 | 43 | 4.967 |  |
| 8 | 4b | 4 | -19 | 49 | 4.333 |  |
| 9 | 4c | 11 | -14 | 49 | 4.289 |  |
| 10 | 5 | -17 | -60 | -19 | 4.851 | 1707 |
| 11 | 5a | -10 | -55 | -16 | 4.651 |  |
| 12 | 5b | -7 | -62 | -22 | 4.217 |  |

**Figure 4.**
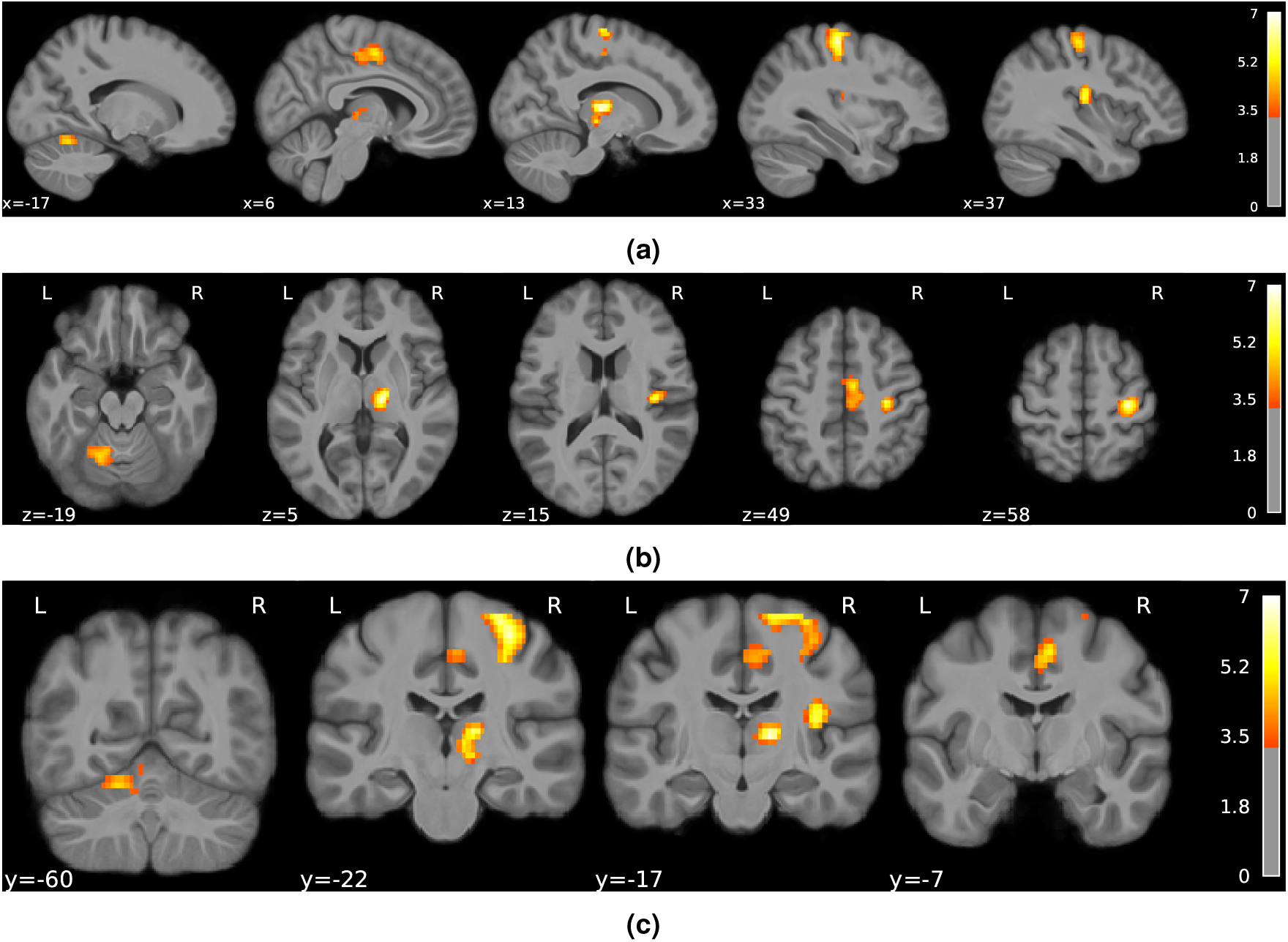
Brain responses to noxious stimuli (high > low pain stimuli) in (a) sagittal, (b) axial and (c) coronal views (colorbar shows Z scores thresholded at z>3.3, FWE corrected p*<*0.05).

Next, we looked at BOLD correlations with the modelled posterior probability of high pain. For any pain stimulus, this reflects the newly calculated probability that the next stimulus will be high, i.e. the dynamic and probabilistic inference of high pain. This analysis identified BOLD responses in the bilateral primary and secondary somatosensory cortex, primary motor cortex and right caudate (fig 5, table 2). We report the opposite contrasts (posterior probability of low pain) in supplementary Figure 3 and Supplementary Table 2.

**Table 2.** Activation clusters associated with the posterior mean p(H) of the Bayesian jump frequency model.

|  | Cluster ID | X | Y | Z | Peak Stat | Cluster Size (mm3) |
| --- | --- | --- | --- | --- | --- | --- |
| 0 | 1 | 66 | -7 | 27 | 6.477 | 4402 |
| 1 | 1a | 52 | -7 | 33 | 5.787 |  |
| 2 | 2 | -62 | -7 | 33 | 5.924 | 2408 |
| 3 | 2a | -46 | -12 | 43 | 4.002 |  |
| 4 | 3 | 21 | -12 | 24 | 4.885 | 1491 |
| 5 | 3a | 11 | -2 | 15 | 4.197 |  |
| 6 | 3b | 13 | -7 | 21 | 4.140 |  |

**Figure 5.**
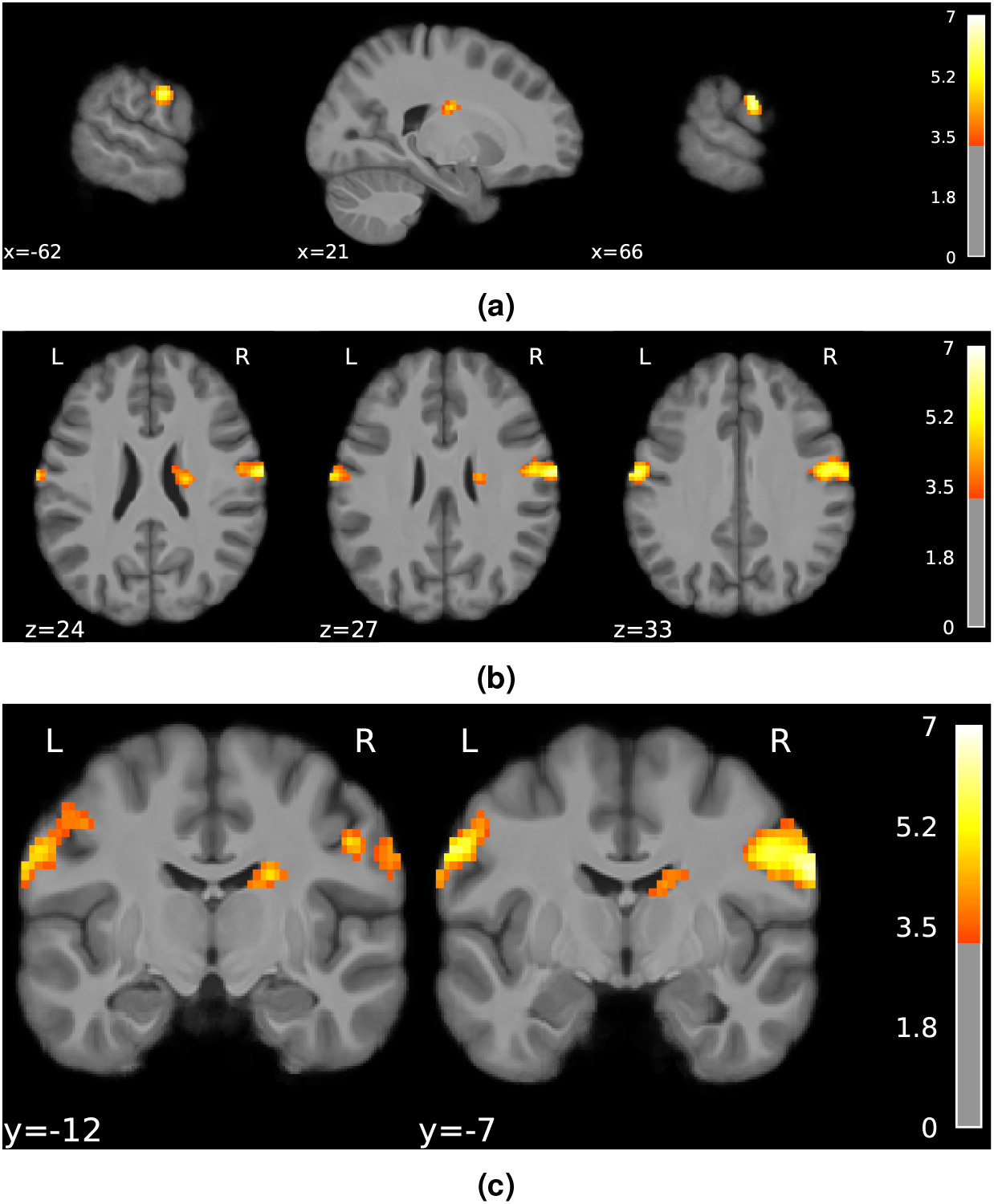
Posterior probability mean of high pain in Bayesian jump frequency model showed activations in the bilateral primary and secondary somatosensory cortex, primary motor cortex and right caudate (FDR corrected p*<*0.001, colorbar shows Z scores > 3.3). (a) sagittal (b) axial and (c) coronal view.

In contrast, a right superior parietal region, bordering with the supramarginal gyrus, was implicated in the computation of the uncertainty (SD) of the posterior probability of high pain, a measure that reflects the uncertainty of pain predictions (figure 6 and table 3). The negative contrast of the posterior SD did not yield any significant cluster.

**Table 3.** Activation clusters associated with the uncertainty of the Bayesian jump frequency model.

|  | Cluster ID | X | Y | Z | Peak Stat | Cluster Size (mm3) |
| --- | --- | --- | --- | --- | --- | --- |
| 0 | 1 | 40 | -48 | 58 | 4.311 | 1186 |
| 1 | 1a | 47 | -38 | 58 | 4.168 |  |
| 2 | 1b | 33 | -41 | 43 | 3.745 |  |
| 3 | 2 | 28 | -58 | 49 | 4.084 | 736 |

**Figure 6.**
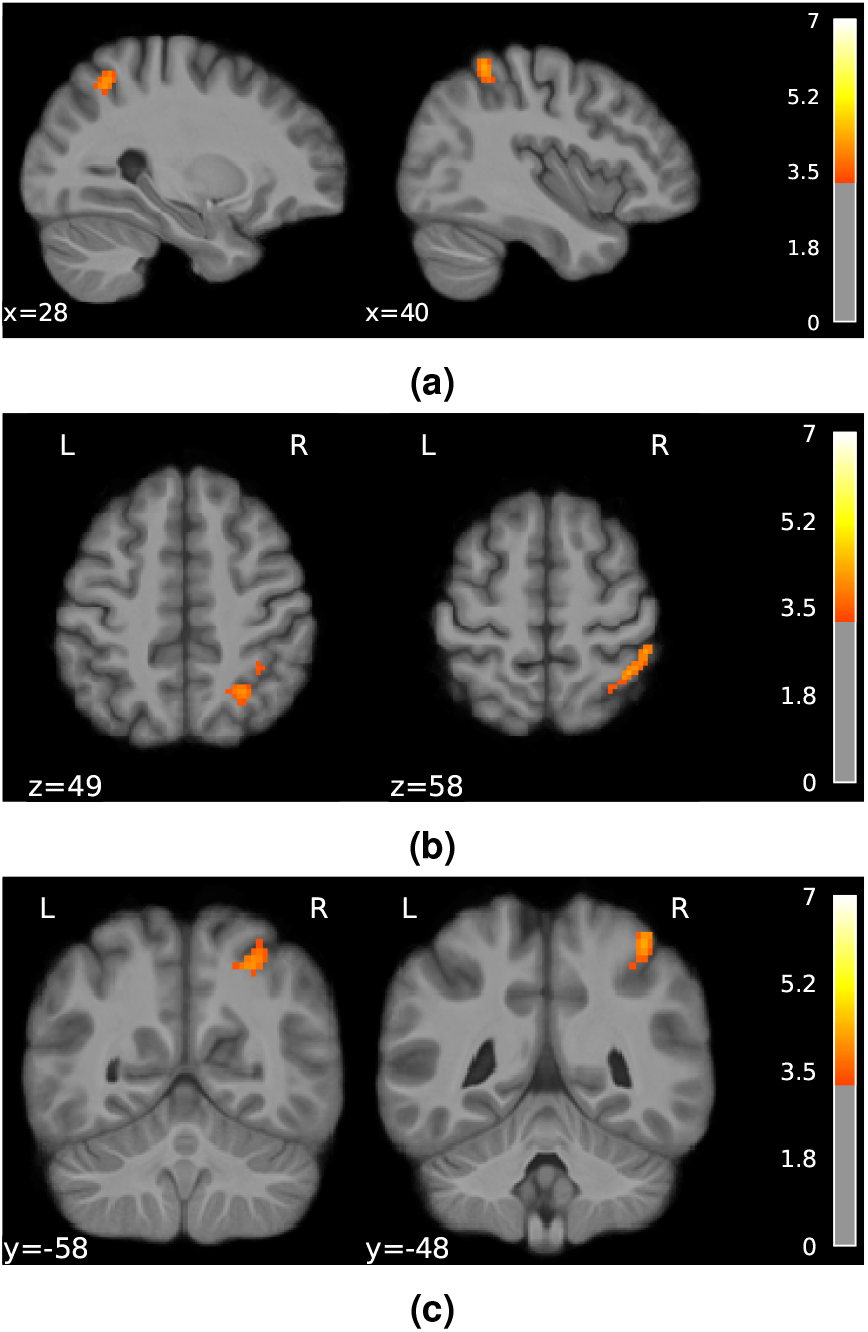
Uncertainty (SD) of the posterior probability of high pain in Bayesian jump frequency model was associated with activations in the right superior parietal cortex (FDR corrected p*<*0.001, colorbar shows Z scores > 3.3). (a) sagittal (b) axial and (c) coronal view.

A key aspect of the Bayesian model is that it provides a metric of the model update, quantified as the KL divergence between successive trial’s posterior distribution. The KL divergence increases when the two successive posteriors are more different from each other, and the opposite when the posteriors are similar. We found that the KL divergence was associated with BOLD responses in left premotor cortex, bilateral dorsolateral prefrontal cortex, superior parietal lobe, supramarginal gyrus, and left somatosensory cortex (fig 7, table 4). For completeness, we report the negative contrast in Supplementary Figure 5 and Supplementary Table 3. Figure 8 overlays the posterior probability with its uncertainty and update (KL divergence). This shows that the temporal prediction of high pain and its update activate distinct, although neighbouring regions in the sensorimotor and premotor cortex, bilaterally. In contrast, the uncertainty of pain predictions activates a right superior parietal region that partially overlaps with the neural correlates of model update.

**Table 4.**
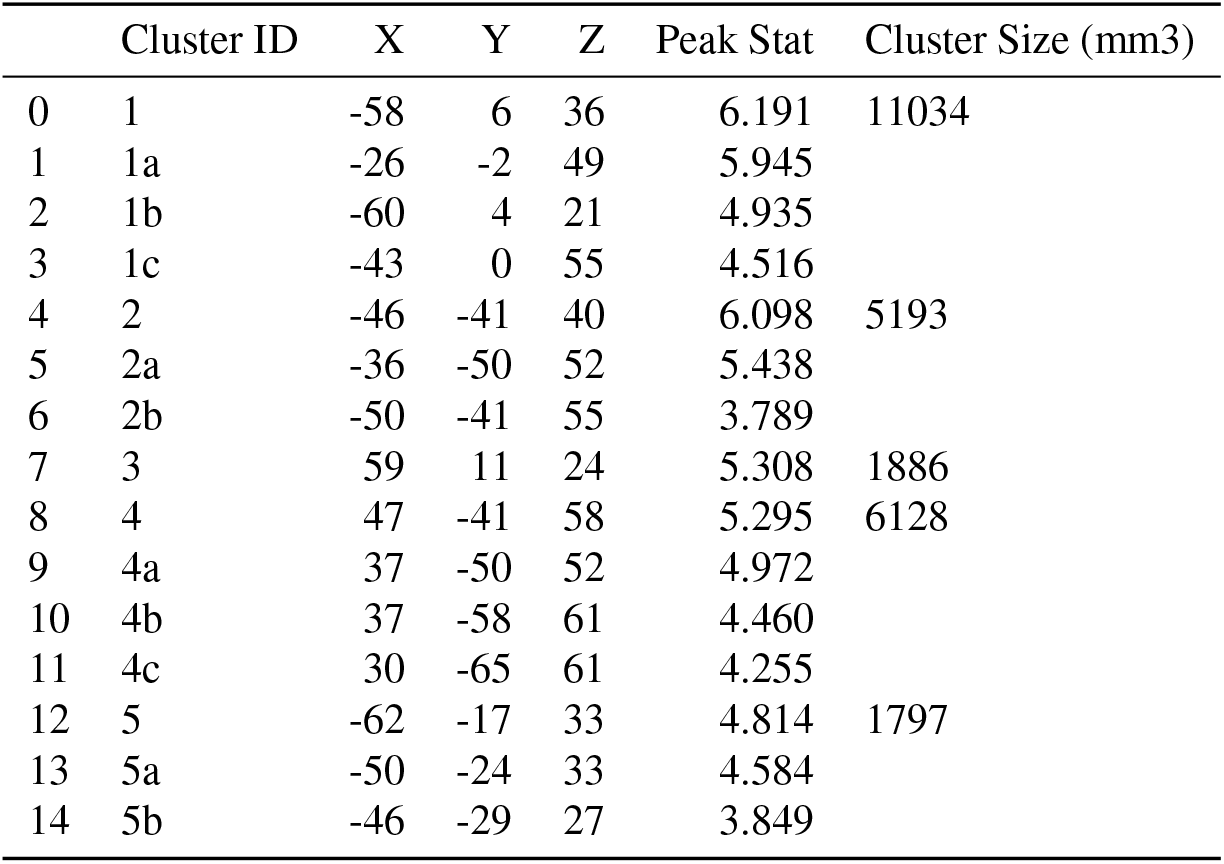
Activation clusters positively associated with the update (KL divergence) of the Bayesian jump frequency model.

|  | Cluster ID | X | Y | Z | Peak Stat | Cluster Size (mm3) |
| --- | --- | --- | --- | --- | --- | --- |
| 0 | 1 | -58 | 6 | 36 | 6.191 | 11034 |
| 1 | 1a | -26 | -2 | 49 | 5.945 |  |
| 2 | 1b | -60 | 4 | 21 | 4.935 |  |
| 3 | 1c | -43 | 0 | 55 | 4.516 |  |
| 4 | 2 | -46 | -41 | 40 | 6.098 | 5193 |
| 5 | 2a | -36 | -50 | 52 | 5.438 |  |
| 6 | 2b | -50 | -41 | 55 | 3.789 |  |
| 7 | 3 | 59 | 11 | 24 | 5.308 | 1886 |
| 8 | 4 | 47 | -41 | 58 | 5.295 | 6128 |
| 9 | 4a | 37 | -50 | 52 | 4.972 |  |
| 10 | 4b | 37 | -58 | 61 | 4.460 |  |
| 11 | 4c | 30 | -65 | 61 | 4.255 |  |
| 12 | 5 | -62 | -17 | 33 | 4.814 | 1797 |
| 13 | 5a | -50 | -24 | 33 | 4.584 |  |
| 14 | 5b | -46 | -29 | 27 | 3.849 |  |

**Figure 7.**
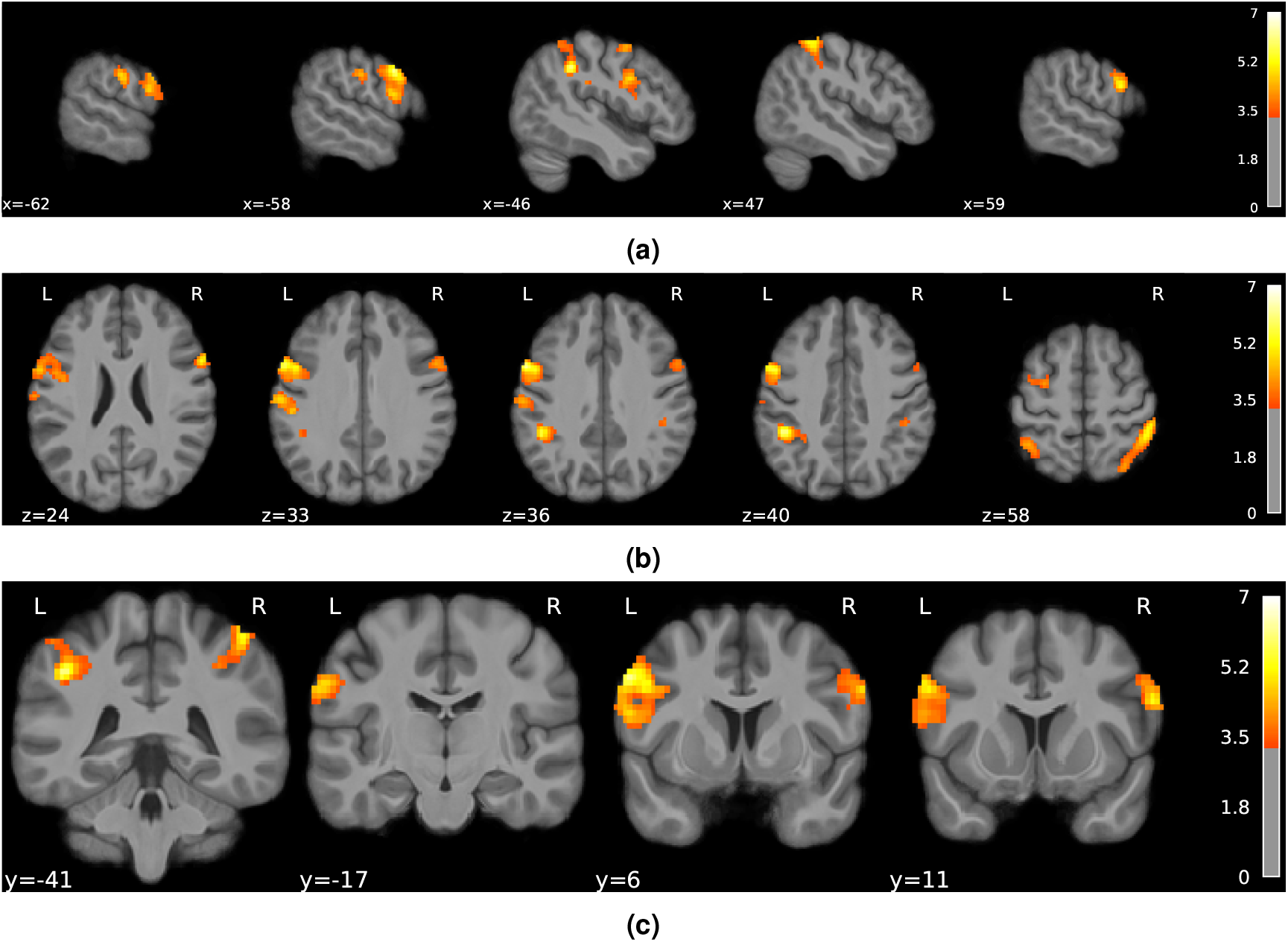
Neural activity associated with the model update, i.e. the KL divergence between posteriors from successive trials (positive contrast), in (a) sagittal, (b) coronal, and (c) axial views (FDR corrected p*<*0.001, colorbar shows Z scores >3.3).

**Figure 8.**
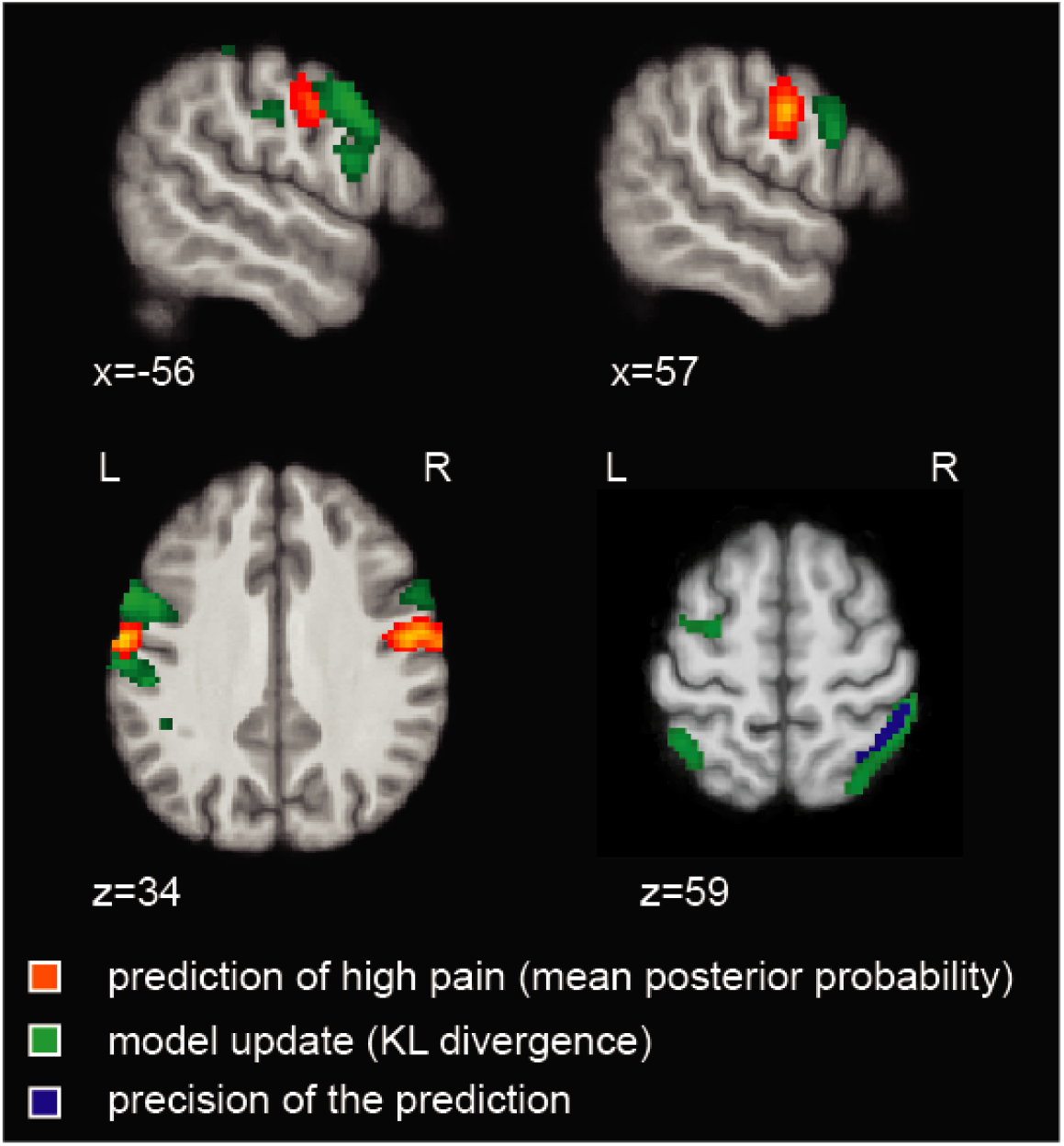
Overlaying the temporal prediction of high pain (mean posterior probability, red-yellow), its uncertainty (SD posterior probability, blue) and the model update (KL divergence between successive posterior distributions, green); (FDR corrected p*<*0.001, colorbar shows Z scores >3.3)

## DISCUSSION

Pain is typically uncertain, and this is most often true when pain persists after injury. When pain persists, the brain needs to be able to track changes in intensity and patterns over time, in order to predict what will happen next and what to do about it. Here we investigated whether, in absence of external cues, the human brain can generate *explicit* (conscious) predictions about the likelihood of forthcoming pain, as these are central to the generation of internal models of pain and can be formally compared to normative models of statistical learning (Dehaene et al., 2015; Meyniel et al., 2016). This study provides evidence that humans can learn and predict the background temporal statistics of pain using optimal

Bayesian inference with dynamic update of beliefs, allowing explicit prediction of the probability of forthcoming pain at any moment in time. Using neuroimaging, we reveal the neural correlates of the internal models of pain predictions. We found distinct neural correlates for the probabilistic, predictive inference of pain and its update. Pain predictions (i.e. mean posterior probability) are encoded in the bilateral, primary somatosensory and motor regions, secondary somatosensory cortex and right caudate, whereas the signal representing the update of the probabilistic model localises in adjacent premotor and superior parietal cortex. The superior parietal cortex is also implicated in the computation of the uncertainty of the probabilistic inference of pain. Overall, the results show that cortical regions typically associated with the sensory processing of pain (primary and secondary somatosensory cortices) encode how likely different pain intensities are to occur at any moment in time, in the absence of any other cues or information; the uncertainty of this inference is encoded in superior parietal cortex and used by a network of parietal-prefrontal regions to update the temporal statistical representation of pain intensity.

The ability of the brain to extract regularities from temporal sequences is well-documented in other sensory domains such as vision and audition (Kourtzi and Welchman, 2019; Dehaene et al., 2015), but pain is a fundamentally different system with intrinsic motivational value and direct impact on the state of the body (Baliki and Apkarian, 2015; Fields, 2018; Seymour, 2019). Despite this fundamental difference, we show that temporal inferences of pain are generated using optimal Bayesian inference - tracking the frequency of low and high intensity pain states and their volatility (i.e. how likely they are to change) based on past experience. A more complex strategy involves trying to infer higher level statistical patterns *within* these sequences, namely representing all the transition probabilities between different states (Meyniel et al., 2016). Although this model fits 1/3 of our subjects best, overall it was not favoured over the simpler frequency learning model, which best describes the behaviour of approximately 2/3 of our sample (figure 3). At this stage it is not clear whether this is because of stable inter-individual differences, or whether given more time, more participants would be able to learn specific transition probabilities. However, it is worth noting that stable, individual differences in learning strategy have been previously reported in visual statistical learning (Karlaftis et al., 2019; Wang et al., 2017).

The Bayesian frequency model is consistent with many other tasks that involve cognitive model learning or acquisition of explicit contingency knowledge across modalities, including pain (Yoshida et al., 2013; Jepma et al., 2018; Hoskin et al., 2019). This reflects a fundamentally different process to pain *response* learning - either in Pavlovian conditioning where simple autonomic, physiological or motoric responses are acquired, or basic stimulus-response (instrumental / operant) avoidance or escape response learning. These behaviours are usually best captured by reinforcement learning models such as temporal difference learning (Seymour, 2019), and reflect a computationally different process (Carter et al., 2006). Having said that, such error-driven learning models have been applied to statistical learning paradigms in other domains before (Orpella et al., 2021), and so here we were able to directly demonstrate that it provided a less accurate model than Bayesian models (figure 3). In contrast to simple reinforcement learning models, Bayesian models allow building an internal, hierarchical model of the temporal statistics of the environment that can support a range of cognitive functions (Honey et al., 2012; Meyniel et al., 2016; Weiss et al., 2021).

A key benefit of the computational approach is that it allows us to accurately map underlying operations of pain information processing to their neural substrates. Our study shows that the probabilistic inference of high pain frequency is encoded in the bilateral sensorimotor cortex, secondary somatosensory cortex, and right caudate (figure 5). The neural correlates of pain predictions arising from predictive Bayesian inference seem to contrast to a certain extent with those arising from value-based learning, which is typically characterised by non-probabilistic model-free learning and involves insula, anterior cingulate and ventromedial prefrontal cortices (Seymour and Mancini, 2020). An exception to this is the observation that the caudate nucleus correlates well with the posterior probability of high pain (i.e. its temporal inference). Although it is difficult to interpret this without an accompanying experimentally-matched value learning task, and without measuring conditioned responses such as autonomic responses, it may represent the parallel or integrative role of caudate in multiple divergent learning processes.

A specific facet of the Bayesian model is the representation of an uncertainty signal, i.e. the posterior SD, and a model update signal, defined as the statistical KL divergence between consecutive posterior distributions. This captures the extent to which a model is updated when an incoming pain stimulus deviates from that expected, taking into account the uncertainty inherent in the original prediction. In our task, the uncertainty of the prediction was encoded in a right superior parietal region, which partially overlapped with a wider parietal region associated with the encoding of the model update (figures 6, 8). This emphasises the close relationship between uncertainty and learning in Bayesian inference (Koblinger et al., 2021). A previous study on statistical learning in other sensory domains reported that a more posterior, intraparietal region, was associated with the precision of the temporal inference (Meyniel and Dehaene, 2017). The role of the superior parietal cortex in uncertainty representation is also evident in other memory-based decision-making tasks, as the superior parietal cortex is more active for low vs. high confidence judgements (Hutchinson et al., 2014; Moritz et al., 2006; Sestieri et al., 2010). In addition to the parietal cortex, the model update signal was encoded in the left premotor cortex and bilateral dorsolateral prefrontal cortex (figure 7), neighbouring regions activated by pain statistical inferences (figure 8). This is particularly interesting, as the premotor cortex sits along a hierarchy of reciprocally and highly interconnected regions within the sensorimotor cortex. The premotor cortex has also been implicated in the computation of an update signal in visual and auditory statistical learning tasks (Meyniel and Dehaene, 2017).

In conclusion, our study demonstrates that the pain system generates probabilistic predictions about the background temporal statistics of pain states, in absence of external cues and using Bayesian-like inference strategy. This extends both current anatomical and functional concepts of what is conventionally considered a ‘sensory pain pathway’, to include encoding not just stimulus intensity (Segerdahl et al., 2015; Wager et al., 2013) and location (Mancini et al., 2012), but the generation of more sophisticated and dynamic internal models of temporal statistics of pain intensity levels. Future studies will need to determine whether temporal statistical predictions modulate pain perception, similarly to other kinds of pain expectations (Büchel et al., 2014; Wiech, 2016; Wager et al., 2004). More broadly, temporal statistical learning is likely to be most important after injury, when continuous streams of fluctuating pain signals ascend nociceptive afferents to the brain, and their underlying pattern may hold important clues as to the nature of the injury, its future evolution, and its broader semantic meaning in terms of the survival and prospects of the individual. It is therefore possible that the underlying computational process might go awry in certain instances of chronic pain, especially when instrumental actions can be performed that might influence the pattern of pain intensity (Jepma et al., 2018; Jung et al., 2017). Thus, future studies could explore both how temporal statistical learning interacts with pain perception and controllability, as well as its application to clinical pain.

## METHODS

### Code and data availability

Raw functional imaging data is deposited at OpenNEURO https://openneuro.org/datasets/ds003836 and derived statistical maps are available at NeuroVault ([upon acceptance]). Sequence generation, task instructions and behavioural data can be found at https://github.com/NoxLab-cam/pain_statistics_3tfrmri. Analysis code can be found at https://github.com/syzhang/tsl_paper.

### Participants

Thirty-five healthy participants (17 females; mean age 27.4 years old; age range 18-45 years) took part in two experimental sessions, 2-3 days apart: a pain-tuning and training session and an MRI session. Each participant gave informed consent according to procedures approved by University of Cambridge ethics committee (PRE.2018.046).

### Protocol

The electrical stimuli were generated using a DS5 isolated bipolar current stimulator (Digitimer), delivered to surface electrodes placed on the index and middle fingers of the left hand. All participants underwent a standardised intensity work-up procedure at the start of each testing day, in order to match subjective pain levels across sessions to a low-intensity level (just above pain detection threshold) and a high-intensity level that was reported to be painful but bearable (>4 out of 10 on a VAS ranging from 0 [‘no pain’] to 10 [‘worst imaginable pain’]). The pain delivery setup was identical for lab-based and MR sessions. After identifying appropriate intensity levels, we checked that discrimination accuracy was >95% in a short sequence of 20 randomised stimuli. This was done to unsure that uncertainty in the sequence task would derive from the temporal order of the stimuli rather than their current intensity level or discriminability. If needed, we tweaked the stimulus intensities to achieve our target discriminability. Next, we gave the task instructions to each participants (openly available https://github.com/NoxLab-cam/pain_statistics_3tfrmri).

After receiving a shock on trial t, subjects were asked to predict the probability of receiving a stimulus of the same or different intensity on the upcoming trial (trial t+1). We informed participants that in the task they “would receive two kinds of stimuli, a low intensity shock and a high intensity shock. The L and H stimuli would be presented in a sequence, in an order set by the computer. After each stimulus, the following stimulus could be either the same or different than the previous one. The computer sets the probability that after a given stimulus (for example L) there would be either L or H” (we showed a visual representation of this example). We asked participants to “always try to guess the probability that after each stimulus there will the same or a different one” and we informed them that “the computer sometimes changes its settings and sets new probabilities”, so to pay attention all the time. We also told them the sequence would be paused occasionally in order to collect probability estimates from participants using the scale depicted in Fig 1. A white fixation cross was displayed on a dark screen throughout the trial, except when a response was requested every 12-18 trials. The interstimulus interval was 2.8-3 seconds. There were 300 stimuli in each block, lasting approx. 8 minutes. Average intensity ratings for each stimulus level were collected after each block during a short break. Low intensity stimuli were felt by participants as barely painful, rated on average 1.39 (SD 0.77) on a scale ranging from 0 (no pain) to 10 (worst pain imaginable). In contrast, high intensity stimuli were rated as more than 4 times higher than low intensity stimuli (mean 5.74, SD 4.85). Participants were given 4 blocks of practice, 2-3 days prior the imaging sessions, and 5 blocks (1500 stimuli in total) during task fMRI.

The sequence of stimuli was unique and generated as in (Meyniel et al., 2016). L and H stimuli were drawn randomly from a 2×2 transition probability matrix, which remained constant for a number of trials (chunks). The probability of a change was 0.014. Chunks had to be >5 and <200 trials long. In each chunk, transition probabilities were sampled independently and uniformly in the 0.15–0.85 range (in steps of 0.05), with the constraint that at least one of the two transition probabilities must be >/< 0.2 than in the previous chunk. Participants were not informed when the matrix was resampled, and a new chunk started.

Behavioural data analysis were conducted with Python packages pandas (pypi version 1.1.3) and scipy (pypi version 1.5.3). Effect size was calculated as Cohen’s d for t-tests.

### Computational modelling of temporal statistical learning

#### Learning models

The models used in comparison are listed as followed:

#### Random (baseline model)

Probabilities are assumed fixed and reciprocal for high and low stimuli, where *p*_*h*_ = 1 − *p*_*l*_ (*p*_*l*_ as free parameter). Uncertainty are also assumed fixed for high/low pain.

#### Rescorla-Wagner (RW model)

Rated probabilities are assumed to be state values, which were updated as

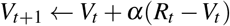

where *R*_*t*_ =1 if stimulus was low, and 0 otherwise. *α* was fitted as free parameter (see (Rescorla et al., 1972)).

#### Bayesian models

Bayesian models update each trial with stimulus identity information to obtain upcoming trial probability from posterior distribution (Meyniel et al., 2016). Using Bayes’ rule, the model parameters *θ*_*t*_ is estimated at each trial *t* provided previous observations *y*_1:*t*_ (sequence of high or low pain), given a model *M*.

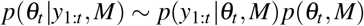

Stimulus information can either be frequency or transition of the binary sequence. There are ‘fixed’ models that assume no sudden jump in stimuli probabilities, and ‘jump’ models that assume the opposite. The four combinations were fitted and compared.

#### 1. Fixed frequency model

For fixed models, the likelihood of parameters *θ* follows a Beta distribution with parameters *N*_*h*_ + 1 and *N*_*l*_ + 1, where *N*_*h*_ and *N*_*l*_ are the numbers of high and low pain in the sequence *y*_1:*t*_. Given that the prior is also a flat Beta distribution with parameters [1,1], the posterior can be analytically obtained with:

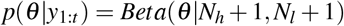

The likelihood of a sequence *y*_1:*t*_ given model parameters *θ* can be calculated as:

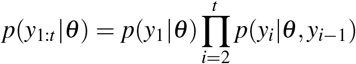

Finally, the posterior probability of a stimulus occurring in the next trial can be estimated with Bayes’ rule:

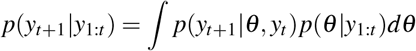

Priors *window* and *decay* were fitted as free parameters, where *window* is the previous *n* trials where frequency of stimuli were estimated, and *decay* is the previous *n* trials where the frequency of stimuli further from current trial were discounted following an exponential decay.

When *window*= *w* is applied, then *N*_*h*_ and *N*_*l*_ are counted within the window of *w* trials *y*_*t*−*w,t*_. When *decay*= *d* is applied, an exponential decay factor 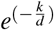 is applied to the *k* trials before their sum is calculated. Both *window* and *decay* were used simultaneously.

#### 2. Fixed transition model

Priors *window* and *decay* were fitted as free parameters as Fixed frequency model above, however, the transition probability was estimated instead of frequency. The likelihood of a stimuli now depends on the estimated transition probability vector *θ* ∼ [*θ*_*h*|*l*_, *θ*_*l*|*h*_] and the previous stimulus pairs *N* ∼ [*N*_*h*| *l*_, *N*_*l*_|_*h*_]. Given that both likelihood and prior can be represented using Beta distributions as before, the posterior result can be analytically obtained as:

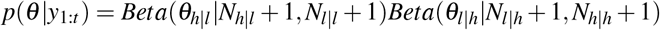

#### 3. Jump frequency model

In jump models, parameter *θ* is no longer fixed, instead it can change from one trial to another with a probability of *p*_*jump*_. Prior *p*_*jump*_ was fitted as a free parameter, representing the subject’s assumed probability of a jump occurring during the sequence of stimuli (e.g. a high *p*_*jump*_ assumes the sequence can reverse quickly from a low pain majority to a high pain majority). The model can be approximated as a Hidden Markov Model (HMM) in order to compute the joint distribution of *θ* and observed stimuli iteratively,

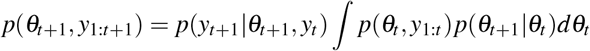

where the integral term captures the change in *θ* from one observation *t* to the next *t* + 1, with probability (1 − *p*_*jump*_) of staying the same and probability *p*_*jump*_ of changing. This integral can be calculated numerically within a discretised grid. The posterior probability of a stimulus occurring in the next trial can then be calculated using Bayes’ rule as

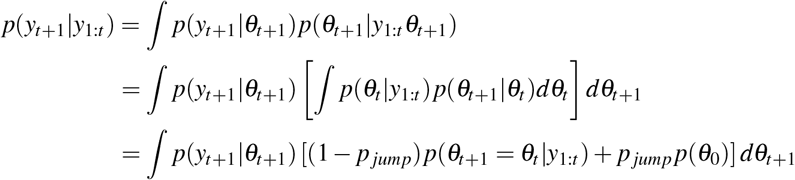

#### 4. Jump transition model

Similar to jump frequency model above, prior *p*_*jump*_ was fitted as a free parameter, but estimating transition instead of frequency. The difference is the stimulus at trial *y*_*t*+1_ now dependent of stimulus at the previous trial, hence the addition of the term *y*_*t*_ in the joint distribution term, shown below.

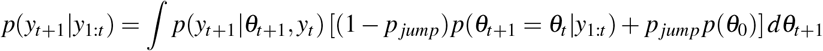

#### KL divergence

Kullback-Leibler (KL) divergence quantifies the distance between two probability distributions. In the current context, it measures the difference between the posterior probability distributions of successive trials. It is calculated as

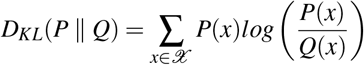

where *P* and *Q* represents the two discrete posterior probability distributions calculated in discretised grids *𝒳*. KL divergence can be used to represent information gains when updating after successive trials (Meyniel and Dehaene, 2017).

#### Subject rated probability

For each individual subject, model predicted probabilities *p*_*k*_ from the trial *k* was used as predictors in the regression:

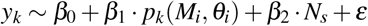

where *y*_*k*_ is the subject rated probabilities, *M*_*i*_ is the *i*th candidate model, *N*_*s*_ is the session number within subject, *β*_0_, *β*_1_, *β*_2_ and *θ*_*i*_ are free parameters to be fitted, and *ε* is normally distributed noise added to avoid fitting errors (Maheu et al., 2019).

#### Model fitting

To estimate the model free parameters from data, Bayesian information criteria (BIC) values were calculated as:

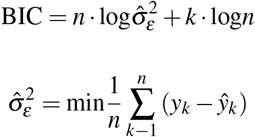

where 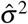 is the squared residual from the linear model above that relates subject ratings to model predicted probabilities, and *n* is the number of free parameters fitted.

We use *fmincon* in MATLAB to minimise the BIC (as approximate for negative log likelihood, Maheu et al. (2019)) for each subject/model. The procedure was repeated 100 times with different parameter initialisation, and the mean results of these repetitions were taken as the fitted parameters and minimised log likelihoods.

#### Model comparison

In general, the best fit model was defined as the candidate model with the lowest averaged BIC. We conducted a random effect analysis with VBA toolbox (Daunizeau et al., 2014), where fitted log likelihoods from each subject/model pair was used as model evidence. With this approach, model was treated as random effects that could differ between individuals. This comparison produces model frequency (how often a given model is used by individuals), model exceedance probability (how likely it is that any given model is more frequent than all other models in the comparison set), and protected exceedance probability (corrected exceedance probability for observations due to chance) (Stephan et al., 2009; Rigoux et al., 2014). These values are correlated and would be considered together when selecting the best fit model.

### Neuroimaging data

#### Data acquisition

First, we collected a T1-weighted MPRAGE structural scan (voxel size 1 mm isotropic) on a 3T Siemens Magnetom Skyra (Siemens Healthcare), equipped with a 32-channel head coil (Wolfson Brain Imaging Centre, Cambridge). Then we collected 5 task fMRI sessions of 246 volumes using a gradient echo planar imaging (EPI) sequence (TR = 2000 ms, TE = 23 ms, flip angle = 78^°^, slices per volume = 31, Grappa 2, voxel size 2.4 mm isotropic, A>P phase-encoding; this included four dummy volumes, in addition to those pre-discarded by the scanner). In order to correct for inhomogeneities in the static magnetic field, we imaged 4 volumes using an EPI sequence identical to that used in task fMRI, inverted in the posterior-to-anterior phase encoding direction. Full sequence metadata are available at OpenNeuro (https://openneuro.org/datasets/ds003836).

#### Preprocessing

Imaging data were preprocessed using fmriprep (pypi version: 20.1.1, RRID:SCR_016216) with Freesurfer option disabled, within its Docker container. Processed functional images had first four dummy scans removed, and then smoothed in an 8mm Gaussian filter in SPM12.

#### GLM analysis

Nipype (pypi version: 1.5.1) was used for all fMRI processing and analysis within its published Docker container. Nipype is a python package that wraps around fMRI analysis tools including SPM12 and FLS in a Debian environment.

First and second level GLM analyses were conducted using SPM12 through nipype. In all first level analyses, 25 regressors of no interest were included from fmriprep confounds output: CSF, white matter, global signal, dvars, std_dvars, framewise displacement, rmsd, 6 a_comp_cor with corresponding cosine components, translation in 3 axis and rotation in 3 axis. Sessions within subject are not concatenated.

In second level analyses, all first level contrasts were entered into a one-sample T-test, with group subject mask applied. The default FDR threshold used was 0.001 (set in Nipype threshold node height_threshold=0.001).

For visualisation and cluster statistics extraction, nilearn (pypi version: 1.6.1) was used. A cluster extent of 10 voxels was applied. Visualised slice coordinates were chosen based on cluster peaks identified. Activation clusters were overlayed on top of a subject averaged anatomical scan normalised to MNI152 space as output by fmriprep.

#### GLM design

All imaging results were obtained from a single GLM model. We investigated neural correlates using the winning Bayesian jump frequency model. All model predictors were generated with the group mean fitted parameters in order to minimise noise. First level regressors include the onset times for all trials, high pain trials, and low pain trials (duration=0). The all trial regressor was parametrically modulated by model-predicted posterior mean of high pain, the KL divergence between successive posterior distributions on jump probability, and the posterior SD of high pain.

For second level analysis, both positive and negative T-contrasts were obtained for posterior mean, KL divergence and uncertainty parametric modulators, across all the first level contrast images from all subjects. A group mean brain mask was applied to exclude activations outside the brain. Given that high and low pain are reciprocal in probabilities, a negative contrast of posterior mean of low pain would be equivalent to the posterior mean of high pain. In addition, high and low pain comparisons were done using a subtracting T-contrast between high and low pain trial regressors. We corrected for multiple comparisons with a cluster-wise FDR threshold of p*<*0.001 for both parametric modulator analyses, reporting only clusters that survived this.

## Supporting information

supplementary results

## ACKNOWLEDGEMENTS

The study was funded by a Medical Research Council Career Development Award to Flavia Mancini (MR/T010614/1) and Wellcome Trust grants to Ben Seynour (097490). We are grateful to Professor Zoe Kourtzi and Dr Michael Lee for helpful discussions about the concept of the study, and to the staff of the Wolfson Brain Imaging Centre for their support during data collection. The authors declare no competing interest.

## AUTHOR CONTRIBUTIONS

FM and BS designed the study. FM collected the data and SZ analysed the data. All authors wrote the paper.

