## supplementary results for "Learning the statistics of pain: computational and neural mechanisms"

### SUPPLEMENTARY MATERIALS AND RESULTS

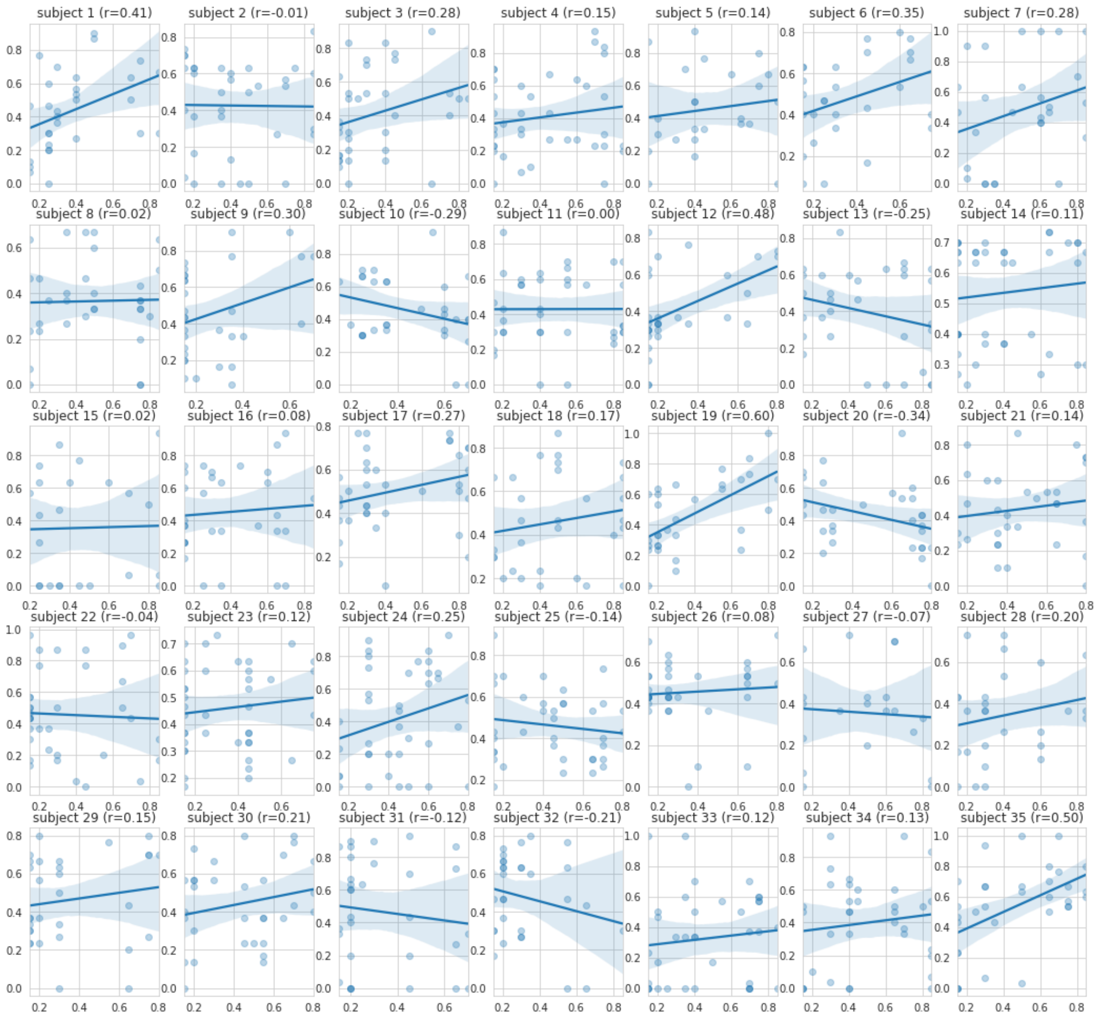


**Figure 1.** Generative vs rated probabilities of p(H|L) for individual participants (mean r=0.117, 74% above 0)

.

| | Cluster ID | X | Y | Z | Peak Stat | Cluster Size (mm3) |
|---|---|---|---|---|---|---|
| 0 | 1 | 47 | -7 | 30 | 7.211 | 6218 |
| 1 | 1a | 61 | -7 | 30 | 7.067 | |
| 2 | 1b | 40 | -26 | 40 | 4.455 | |
| 3 | 1c | 47 | -19 | 43 | 4.453 | |
| 4 | 2 | -60 | -7 | 33 | 6.263 | 12669 |
| 5 | 2a | -67 | -10 | 24 | 5.806 | |
| 6 | 2b | -53 | -10 | 43 | 5.170 | |
| 7 | 2c | -26 | -41 | 49 | 5.074 | |
| 8 | 3 | 4 | -38 | 61 | 5.249 | 2354 |
| 9 | 4 | -22 | -79 | 36 | 5.231 | 1006 |

**Table 1.** Low pain > high pain stimuli activation clusters (FWE p<0.05)

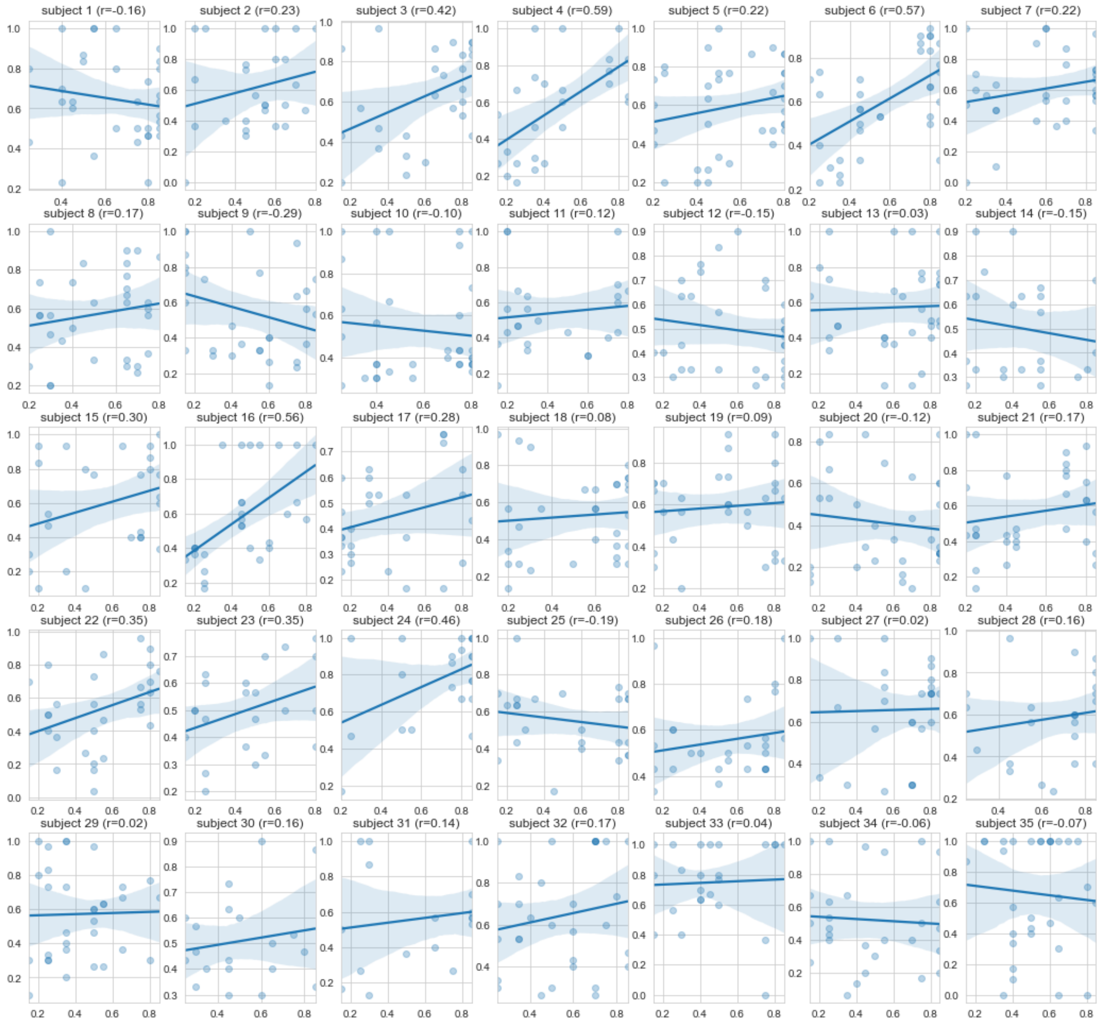


**Figure 2.** Generative vs rated probabilities of p(H|H) for individual participants (mean r=0.117, 74% above 0)

.

| | Cluster ID | X | Y | Z | Peak Stat | Cluster Size (mm3) |
|---|---|---|---|---|---|---|
| 0 | 1 | 28 | -22 | 65 | 6.158 | 5265 |
| 1 | 1a | 30 | -19 | 49 | 6.012 | |
| 2 | 1b | 16 | -12 | 65 | 5.148 | |
| 3 | 1c | 23 | -10 | 68 | 4.575 | |
| 4 | 2 | 37 | -17 | 15 | 6.013 | 1311 |
| 5 | 3 | 13 | -22 | 8 | 5.407 | 1886 |
| 6 | 4 | 11 | -17 | 49 | 4.683 | 2372 |
| 7 | 4a | 0 | -2 | 43 | 4.439 | |

**Table 2.** Bayesian model (jump frequency) posterior mean p(L) activation clusters

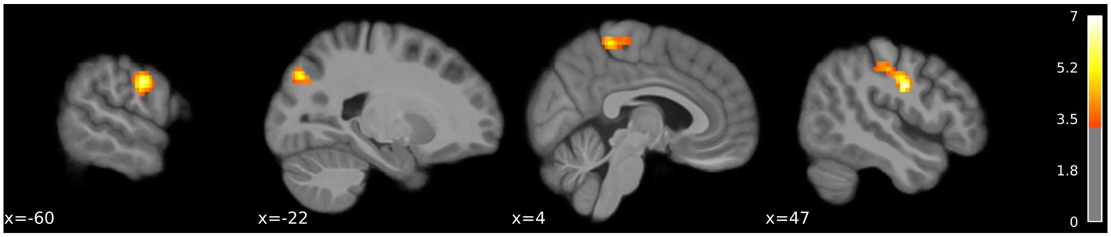


**(a)**

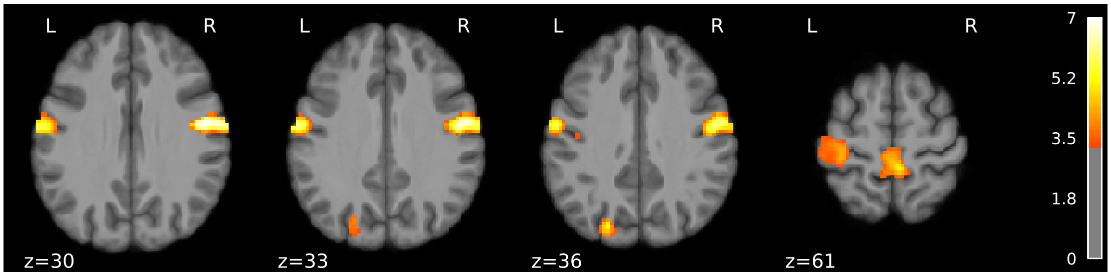


**(b)**

**Figure 3.** Brain responses to noxious stimuli, low>high pain contrast (colorbar shows Z scores, FWE corrected $p<0.05$). (a) sagittal and (b) axial views.

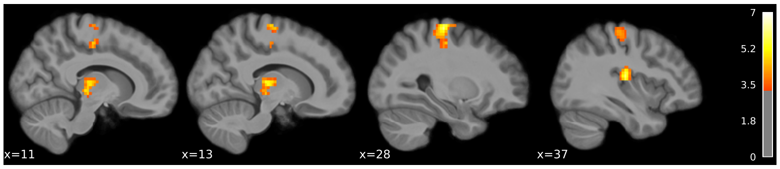


**(a)**

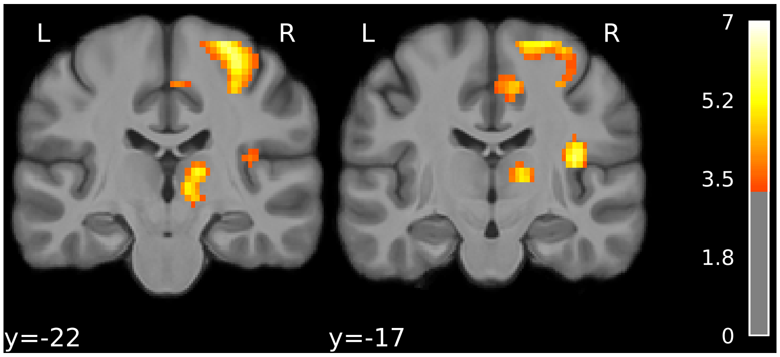


**(b)**

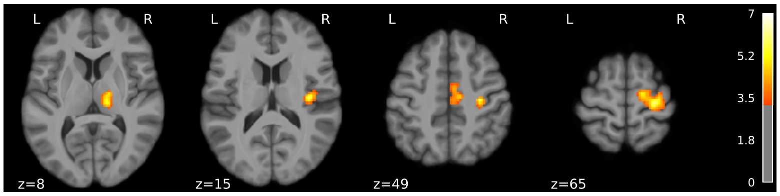


**(c)**

**Figure 4.** Posterior probability mean of low pain in Bayesian jump frequency model (FDR corrected $p<0.001$, colorbar shows Z scores, thresholded at $Z>3.3$). (a) sagittal (b) coronal and (c) axial view.

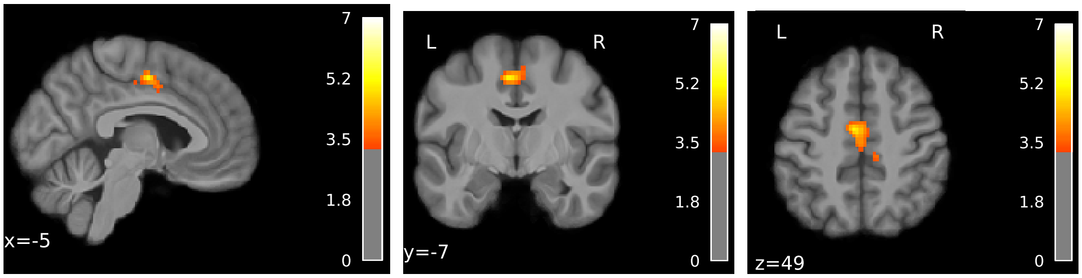


(a) (b) (c)

**Figure 5.** KL divergence / (a-c) Negative contrast (FDR corrected p<0.001, colorbar shows Z scores, thresholded at Z>3.3)

| | Cluster ID | X | Y | Z | Peak Stat | Cluster Size (mm3) |
|---|---|---|---|---|---|---|
| 0 | 1 | -5 | -7 | 49 | 5.096 | 3414 |
| 1 | 1a | 9 | -19 | 43 | 4.766 | |
| 2 | 1b | 0 | -17 | 46 | 4.513 | |

**Table 3.** Bayesian model (jump frequency) KL divergence (negative contrast) activation clusters
